# The neural basis of effort valuation: A meta-analysis of functional magnetic resonance imaging studies

**DOI:** 10.1101/2021.01.08.425909

**Authors:** Paula Lopez-Gamundi, Yuan-Wei Yao, Trevor T-J. Chong, Hauke R. Heekeren, Ernest Mas Herrero, Josep Marco Pallares

**Author notes:** Corresponding Author: Paula Lopez-Gamundi or Yuan-Wei Yao. Equal contribution (Paula Lopez-Gamundi, Yuan-Wei Yao). Equal contribution (Trevor T-J. Chong, Hauke R. Heekeren, Ernest Mas Herrero, Josep Marco Pallares).

## Abstract

Choosing how much effort to expend is a critical for everyday decisions. While effort-based decision-making is altered in common psychopathologies and many neuroimaging studies have been conducted to examine how effort is valued, it remains unclear where the brain processes effort-related costs and integrates them with rewards. Using meta-analyses of combined maps and coordinates of functional magnetic resonance imaging (fMRI) studies (total N = 22), we showed that raw effort demands consistently activated the pre-supplementary motor area (pre-SMA). In contrast, the net value of effortful reward consistently activated regions, such as the ventromedial prefrontal cortex (vmPFC) and ventral striatum (VS), that have been previously implicated in value integration in other cost domains. The opposite activation patterns of the pre-SMA and vmPFC imply a double dissociation of these two regions, in which the pre-SMA is involved in pure effort cost representation and the vmPFC in net value integration. These findings advance our understanding of the neural basis of effort-related valuation and reveal potential brain targets to treat motivation-related disorders.

## 1. Introduction

Every day, we are faced with choices about whether to invest effort to attain certain goals (Bailey et al., 2016; Salamone et al., 2009). These effort demands are often regarded as costly, such that individuals tend to avoid one action if it requires too much effort (Kool et al., 2010; Kurniawan et al., 2010, 2011). The ability to accurately weigh energy requirements against potential benefits (e.g., “effort-based decision-making”), is therefore crucial for optimal goal-directed action, and alterations in this function are believed to be a core component of motivational disorders, such as apathy (Chong and Husain, 2016; Hartmann et al., 2015; Husain and Roiser, 2018), and have been found across a variety of psychopathologies, including depression (Treadway et al., 2012; Yang et al., 2014), schizophrenia (Barch et al., 2014; Park et al., 2017), Parkinson’s disease (Chong, 2018; den Brok et al., 2015; Le Heron et al., 2018), and substance dependence (Grodin et al., 2016). Due to its clear clinical importance, there has been a recent surge of interest in how effort devalues prospective rewards, and such studies have demonstrated that effort might be a unique cost, distinct from other more investigated cost domains, such as risk and delay. However, work on the neural mechanisms underlying effort-based valuation have yielded heterogeneous results, and the question of how humans integrate effort and reward remains a subject of contention.

Most behavioral economic theories of reward-related behavior rely on the assumption that an organism weighs a reward and its associated costs to generate a net value of an option (Kahneman and Tversky, 1979; Sutton and Barto, 1998; Von Neumann and Morgenstern, 1990). A popular hypothesis proposes that, to effectively compare different options, the net value of each must be represented in a ‘common currency’ (Padoa-Schioppa, 2011; Rangel et al., 2008; Westbrook and Braver, 2015). A network of regions, including the ventromedial prefrontal cortex (vmPFC; and adjacent orbitofrontal cortex) and ventral striatum (VS), have been repeatedly implicated in the encoding of the net value of rewards discounted by the costs associated with obtaining them (Bartra et al., 2013; Levy and Glimcher, 2012). Based on these data, this valuation network is posited to be ‘domain-general’, as it tracks net value representations regardless of the nature of the reward (e.g., primary vs secondary) (Bartra et al., 2013; Sescousse et al., 2013) or of the type of cost (e.g., risk vs delay) (Kable and Glimcher, 2007; Peters and Büchel, 2009; Prévost et al., 2010).

However, much of these data have focused on outcome-related costs such as risk or delay. Notably, research on effort-based valuation suggests a limited role for the vmPFC and VS for value integration. Instead, other frontal regions beyond this core valuation network, including the anterior cingulate cortex (ACC), supplementary motor area (SMA), and anterior insula (AI), have been shown to signal net value discounted by effort costs (Arulpragasam et al., 2018; Camille et al., 2011; Chong et al., 2017; Klein-Flugge et al., 2016; Massar et al., 2015; Skvortsova et al., 2014; Walton et al., 2003). These findings are consistent with animal studies showing that lesions to the ACC, but not the nucleus accumbens, prelimbic/infralimbic cortex (homologous to the vmPFC), or orbitofrontal cortex, reduce the amount of effort rats invested for rewards (Rudebeck et al., 2006; Walton et al., 2009, 2003). Furthermore, neural activity in the ACC, as measured by single unit recordings, varies with cost-benefit weighting (Hillman and Bilkey, 2012, 2010) and effort-related choice (Cowen et al., 2012). This body of work thus raises the possibility that a distinct frontal network is specifically recruited to integrate effort-related value.

On the other hand, these frontal regions (i.e. ACC, pre-SMA, AI, etc.) are also commonly implicated in cognitive control processes (Wu et al., 2020), which may overlap or obscure value signals. For example, value-based decision-making may trigger cognitive control functions such as conflict detection and response inhibition (Botvinick and Braver, 2015; Botvinick et al., 2001), surprise and/or prediction error signaling (Vassena et al., 2020, 2017), and invigoration of goal-directed behavior (Kouneiher et al., 2009; Kurniawan et al., 2013; Mulert et al., 2005).Therefore, it is plausible that these regions are recruited to prepare and invigorate behaviors necessary for realizing a prospective reward instead of for computing prepotent net values per se. Another situation that requires cognitive control is difficult decision-making when two simultaneously presented options have similar net value (Chong et al., 2017; Hunt et al., 2012; Klein-Flugge et al., 2016; Massar et al., 2015; Shenhav et al., 2013). Indeed, studies that have independently manipulated net value and decision difficulty showed that these frontal regions, particularly the dorsal ACC, specifically tracked decision difficulty (Hogan et al., 2017; Westbrook et al., 2019) while, in contrast, the vmPFC uniquely tracked net value (Westbrook et al., 2019). Taken together, these findings suggest that this distinct frontal network is recruited more specifically for cognitive control, such as response planning and option comparison, and that effort-related value integration is still processed in the core valuation network (e.g., vmPFC and VS) that have been identified in other cost domains.

The inconsistencies in previous studies may be related to several issues. For example, some may have been statistically underpowered due to small sample sizes, which may have reduced the probability of detecting significant effects, and/or reduce the reliability of their findings (Müller et al., 2018; Poldrack et al., 2017). Furthermore, the specific effort requirements of each task may have induced different patterns of brain activity, making it difficult to judge whether findings from individual studies can be generalized to the cognitive process of interest. A promising approach to address these issues is to quantitatively synthesize fMRI data across multiple studies using an image-based meta-analysis (Muller et al., 2018). Relative to traditional meta-analyses based only on peak coordinates of significant activity, an image-based meta-analytic approach uses the full information of the statistical maps from each study, and has greater power to detect small effect sizes (Luijten et al., 2017; Salimi-Khorshidi et al., 2009). A previous study showed that even the inclusion of 20% of statistical maps for included studies could significantly improve the precision of a meta-analysis (Radua et al., 2012).

Here, we conducted a hybrid coordinate- and image-based fMRI meta-analysis to identify the neural correlates of effort-related cost processing and value integration. Considering their critical roles in response planning, we hypothesized that frontal regions like the ACC, SMA, and AI would be consistently involved in representing prospective effort, independent of the reward offer. We also aimed to test whether effort-related value integration (i.e., the integration of reward value with the effort required to obtain it) relied on the core valuation areas such as the vmPFC and VS or broader frontal regions.

## 2. Materials and Methods

### 2.1 Literature Screen, Data Collection, and Preparation

#### 2.1.1 Exhaustive Literature Search

We conducted a systematic literature search to identify neuroimaging studies on prospective effort and the integration of reward value and effort costs in healthy adults. Candidates for inclusion were initially identified by searching PubMed, ProQuest, and Web of Science on June 29, 2020 using the grouped terms (“fMRI” OR “functional magnetic resonance imaging”) AND (“effort discounting” OR “effort-based decision-making” OR “effort valuation” OR “effort anticipation” OR “cost-benefit valuation” OR “cognitive effort” OR “physical effort” OR “effort expenditure” OR “effort allocation” OR “effortful goal directed action” OR “reward related motivation” OR “reward related effort”). Searches were limited to human studies where databases would allow. 121, 787, and 127 studies were identified on PubMed, ProQuest, and Web of Science, respectively. We also searched existing in-house reference libraries and names of prominent authors in the field, resulting in the addition of candidate studies. 934 candidate studies remained after search results were pooled and duplicates removed. Two researchers (PL-G, Y-WY) then independently reviewed the title and abstract of candidate papers to determine relevance, resulting in a pool of 72 studies that underwent a full-text review (Figure 1).

**Figure 1.**
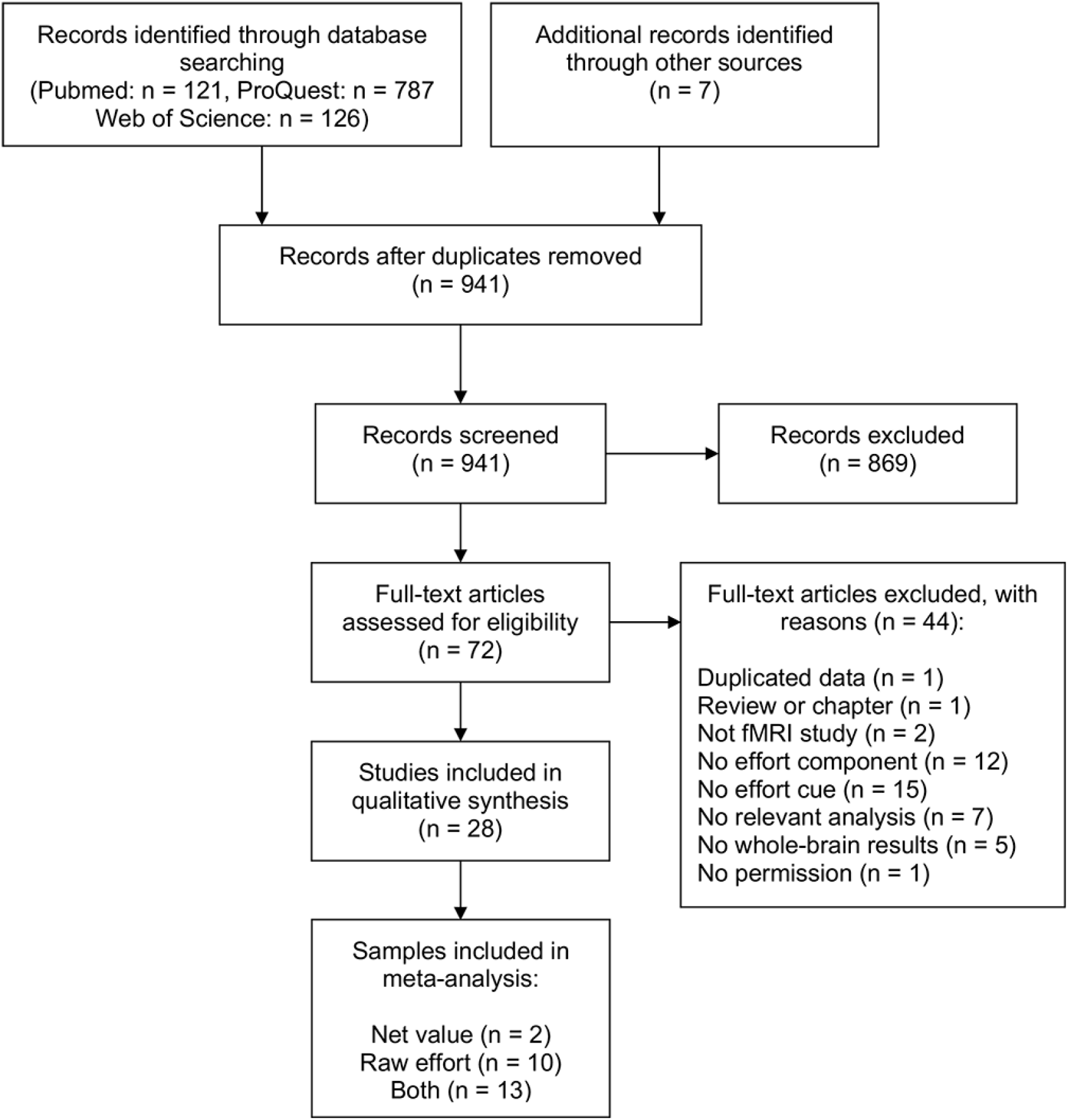
Preferred Reporting Items for Systematic Reviews and Meta-Analyses (PRISMA) flow diagram.

#### 2.1.2 Inclusion/Exclusion Criteria

Studies were included if they: 1) had a healthy adult human sample in the non-elderly age range (ages 18 to 65, with one exception detailed below); 2) used functional MRI; 3) either reported or referenced a whole-brain analysis; and 4) utilized a task with an effort component with clear effort (or combined effort and reward) cues during an ‘anticipation’ phase. Please note that ‘anticipation’ in this case refers to the evaluation of prospective effortful rewards before or during decision-making, and does not include anticipatory responses to reward post-effort exertion (e.g., the ‘evaluation’ phase described in Assadi et al., (2009)).

To ensure that the selected studies could be meaningfully compared, we limited the final corpus to those that used experimental paradigms with certain characteristics. First, because studies have found that loss and gain are asymmetric and partially dissociable (Chen et al., 2020; Porat et al., 2014; Tanaka et al., 2014), we excluded studies that used paradigms with only loss conditions, or that only conducted gain vs loss comparisons. Second, we excluded studies that only used a single speeded response as its effort component (e.g. classical Monetary Incentive Delay task (Knutson et al., 2000)), as this was not deemed as a significant effort demand, and other reviews and meta-analyses focusing on reward anticipation with these paradigms can be found elsewhere (Diekhof et al., 2012; Knutson and Greer, 2008; Wilson et al., 2018). Finally, we only included those studies which measured activity during the *prospective* valuation of an action and its rewards, rather than only at the time of reward outcome, as estimates of previously expended effort can be biased by reward receipt (Pooresmaeili et al., 2015).

We contacted the corresponding authors of 28 candidate studies to request whole-brain statistical maps for the analyses of interest, and received whole-brain statistical maps or peak coordinates from 25 studies. In cases where only between-group (e.g. clinical studies) and/or ROI results were reported, we contacted corresponding authors to inquire about the availability of whole-brain results for relevant contrasts in healthy adult subjects. If images were not available, we requested they provide us with peak activation foci in stereotactic spatial coordinates (i.e., Talairach or MNI space), together with the direction of the effect (positive or negative).

#### 2.1.3 Data collection and preparation

We performed two analyses of interest. The first examined activity related to the raw effort involved in the option itself. We included analyses that examined high vs. low effort demands (i.e., categorical contrasts) and those that examined continuous changes in effort (i.e., parametric modulation). The second analysis examined activity related to the prospective net value of an effortful reward. Whenever possible, we used the contrast related to the net value of a single option (i.e., the subjective value of the chosen option discounted by the effort required to obtain it). When this contrast was unavailable, we used the contrast related to the differences between options instead. Studies that only investigated BOLD activity associated with interactions between reward and effort were excluded, as they did not rely on the same discounting assumptions as other measures of net value. It should be noted that one study (Nagase et al., 2018) included two experiments with six common participants, so we selected the experiment with a larger sample size for the meta-analysis. In another study (Chong et al., 2017), all participants took part in both cognitive and physical effort-based decision-making tasks. Thus, we combined the statistical maps from both tasks to avoid selection bias. Finally, one study (Seaman et al., 2018) had a sample that included participants ranging from 22 to 83 years old. However, the authors of this study provided whole-brain maps that controlled for the effect of age, and we chose to include this data in the net value meta-analysis.

#### 2.1.4 Final Corpus

As shown in Figure 1, 25 studies were ultimately included in the final corpus of studies, which were considered in one or both meta-analyses on raw effort evaluation and effort-reward integration. The raw effort valuation analysis included 15 maps (65%) and 7 coordinates for raw effort processing, resulting in 22 studies with a total sample of N = 549 (mean = 24.95; median = 22.5, range = [16-50]). A description of the final corpus of studies can be found in Table 1. The value integration analysis included 11 maps (73%) and 4 coordinates, resulting in 15 studies, with a total sample of N = 428 participants (mean = 28.5; median = 23, range = [16-75]).

**Table 1.**
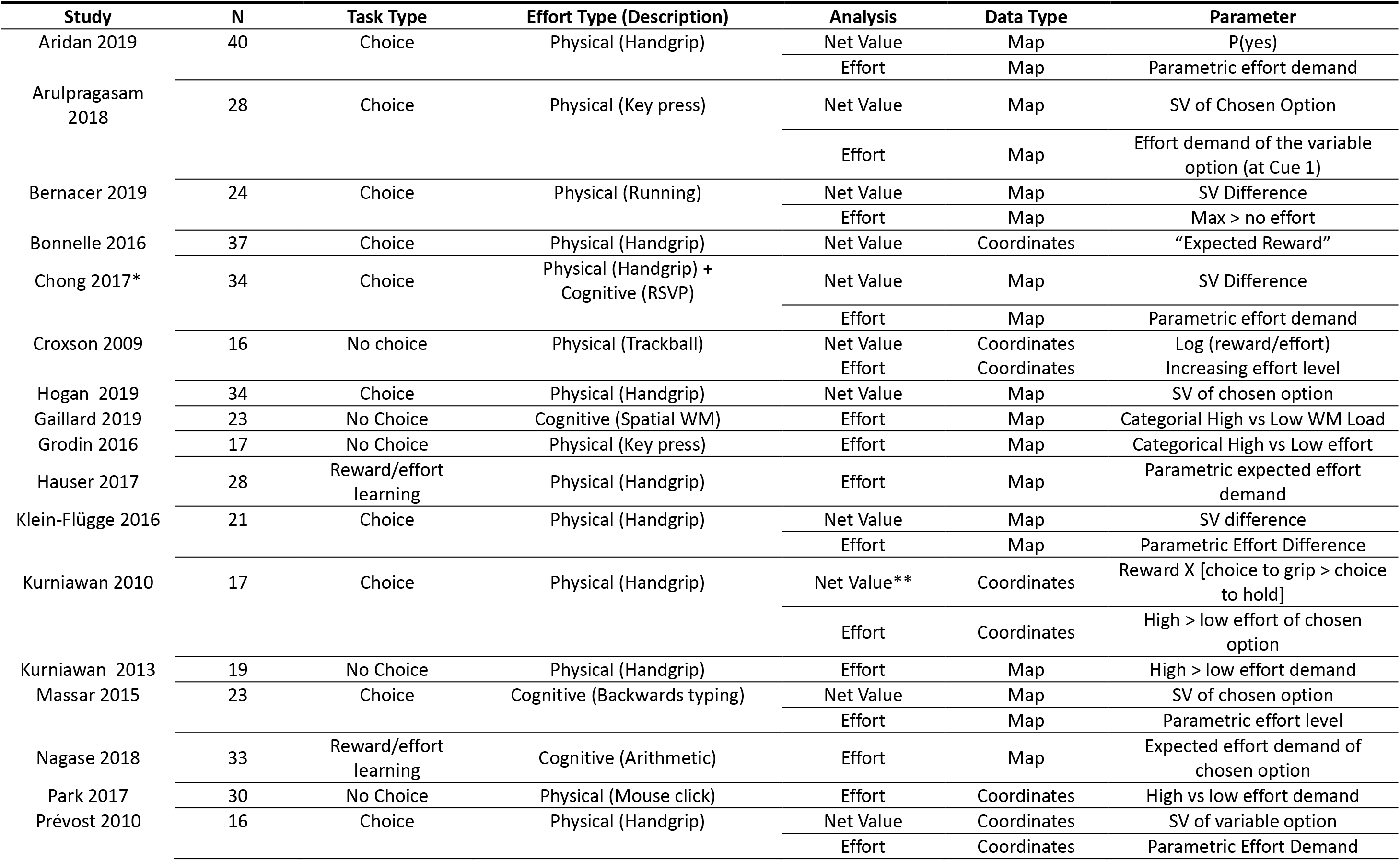
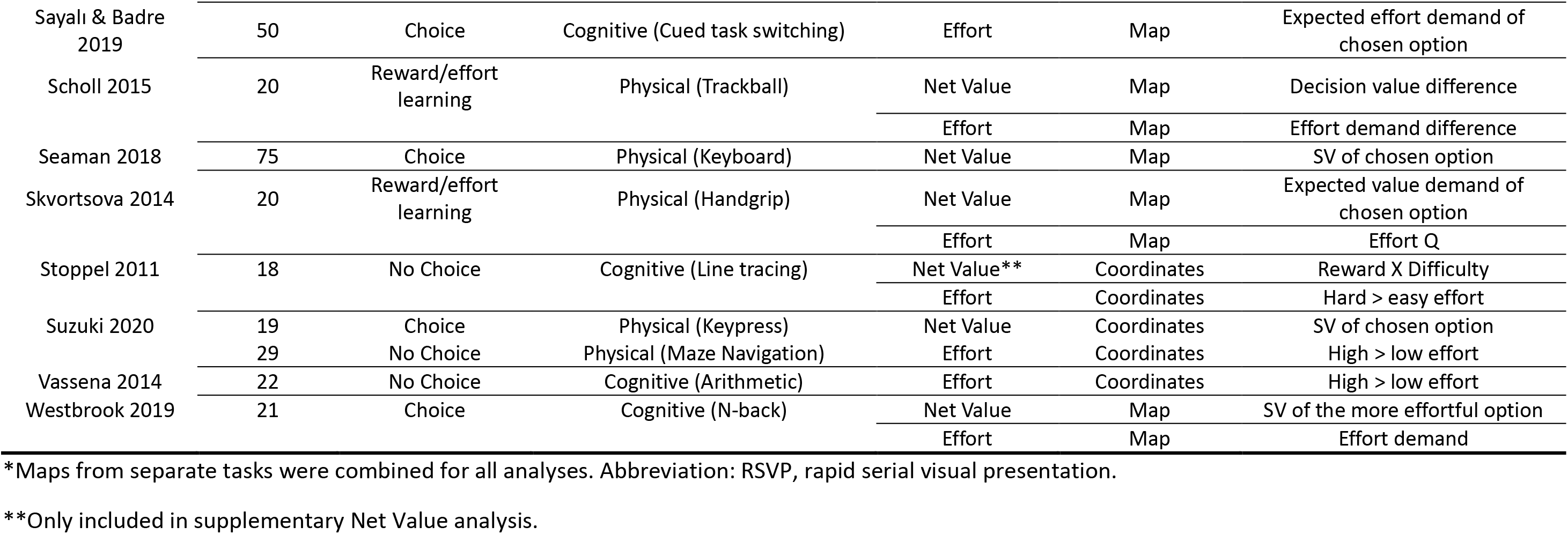
Summary of Included Studies

### 2.2 Meta-Analytic Procedures

#### 2.2.1 Seed-based d Mapping

The combined image- and coordinate-based meta-analyses were performed using the software Seed-based *d* Mapping with Permutation of Subject Images (SDM-PSI, version 6.21; https://www.sdmproject.com). SDM-PSI preserves the information about the sign of the effect and the methods have been validated in previous studies (Albajes-Eizagirre et al., 2019; Radua et al., 2012). During preprocessing, SDM-PSI recreated voxel-level maps of standardized effect sizes (i.e., Hedge’s g) and their variances and allowed the incorporation of both whole-brain t-maps and peak information (i.e., coordinates and t-values). The inclusion of statistical maps can substantially increase the sensitivity of meta-analyses compared with the pure coordinate-based approach (Radua et al., 2012). When *t*-maps were unavailable, SDM-PSI estimated them based on coordinates and their effect sizes using anisotropic kernels (Radua et al., 2014).

#### 2.2.2 Meta-analysis

Two separate whole-brain meta-analyses were conducted to examine consistent neural correlates of prospective effort and net value processing, respectively. Random-effect models were used to assess the mean effect size of each study, where the weight of a study is the inverse of the sum of its variance and the between-study variance. SDM z-maps were generated by dividing the voxel-wise effect sizes by their standard errors. As these z-values may deviate from a normal distribution, a null-distribution was estimated for each meta-analysis from 50 whole-brain permutations.

##### 2.2.2.1 Region-of-Interest (ROI) Analysis

To directly investigate the involvement of key brain regions in effort-related cost processing and value integration, we focused on seven *a priori* regions of interest (ROIs) derived from an independent meta-analysis (Bartra et al., 2013) that examined valuation network in general. Those ROIs included: the vmPFC, right and left VS, ACC, pre-SMA, and right and left AI, which generally covered the core valuation network and additional frontal regions of interest. A spherical mask of radius 6mm was created for each ROI centered on the respective peak coordinates. Mean effect sizes and variances of those ROIs were extracted from individual studies with statistical maps.

##### 2.2.2.2 Whole-Brain Analysis

We also examined the whole-brain results beyond these *a priori* ROIs. To reduce the false-positive results due to multiple comparisons, we applied a familywise error (FWE) correction with 1000 subject-based permutations (Albajes-Eizagirre et al., 2019). In accordance with SDM-PSI’s recommendations, a threshold-free cluster enhancement (TFCE) corrected *p* < 0.025 was used (Albajes-Eizagirre et al., 2019).

In addition, we performed a conjunction analysis to identify regions that were associated with both raw effort demand and net value. For exploratory purposes, we created maps using a voxel-level uncorrected threshold of *p* < 0.001 and a cluster size > 20 voxels for both meta-analyses. Masks were generated from significant clusters that activated or deactivated in both processes (i.e., based on absolute values). We then used SPM12 (http://www.fil.ion.ucl.ac.uk/spm) to perform a conjunction analysis to extract overlapping areas for both processes, regardless of the direction (Culter & Campbell-Meiklejohn, 2019).

Finally, we conducted two supplementary analyses. First, because of the possible role of the dorsal ACC and other frontal regions in signaling choice difficulty, we were interested in assessing if our findings were influenced by studies that used net value differences as the parameter, rather than the net value of the chosen option. Thus, we repeated the meta-analysis with a subgroup of studies that used parameters only representing the net value of a single option (N=11). Second, because net value can also be more broadly defined as an interaction between reward and effort, we repeated the net value meta-analysis by including the coordinates of two additional studies (Kurniawan et al., 2010; Stoppel et al., 2011) that used interaction parameters (e.g. Reward X Effort) as opposed to traditional discounting parameters of net value (e.g. SV). These analyses were conducted using the same procedures described above.

#### 2.2.3 Heterogeneity and Publication Bias

Areas of significant activation were assessed for heterogeneity and publication bias. For each meta-analysis, peaks with heterogeneity *l*^*2*^ values > 20% were flagged and inspected. In order to assess publication bias, Hedge’s *g* effect size estimates were extracted at the study level for peak voxels of significant clusters. Funnel plots were created and visually inspected. Egger regression tests (Egger et al., 1997) were conducted to quantitatively test the asymmetry of each funnel plot.

### 2.3 Data Availability

Unthresholded z-maps of our results are available at NeuroVault: https://neurovault.org/collections/9286/. The TFCE-corrected maps as well as publication bias and heterogeneity data are available from the corresponding authors upon request.

## 3. Results

### 3.1 ROI Analysis

To directly examine the roles of key regions in raw effort prospect and effort-reward integration, we focused on seven *a priori* ROIs. Results are summarized in Table 2. The vmPFC showed consistent activations related to net value and deactivations related to prospective effort. The bilateral VS showed a similar activity pattern, but smaller effect sizes for both analyses. In contrast, the pre-SMA showed consistent activations related to effort demand and deactivations related to net value. The ACC and bilateral AI showed similar activity pattern, but smaller effect sizes for both analyses. Figures 2 and 3 show the Hedge’s g effect sizes for raw effort prospect and net value analyses in the vmPFC and pre-SMA ROIs. The forest plots for other regions were shown in Figure S1-S10.

**Figure 2.**
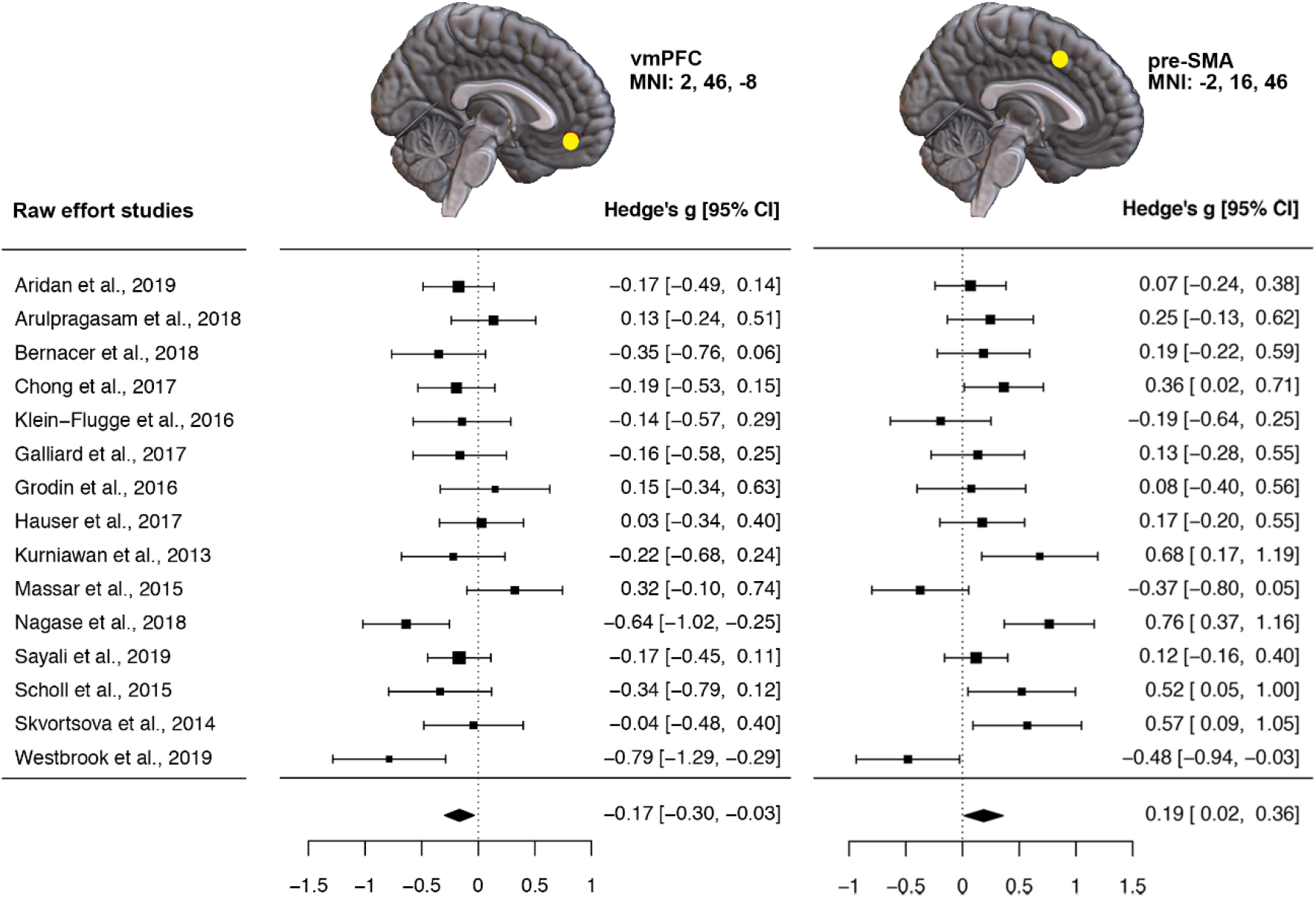
Forest plot illustrating activation related to effort demand in the vmPFC and pre-SMA ROIs in studies with statistical maps. Contrary to our findings for net value signaling, the pre-SMA is activated (Hedge’s g= 0.20, 95% CI [0.02, 0.37]) and the vmPFC is deactivated (Hedge’s g= −0.17, 95% CI [−0.30, - 0.03]) when tracking pure prospective effort.

**Figure 3.**
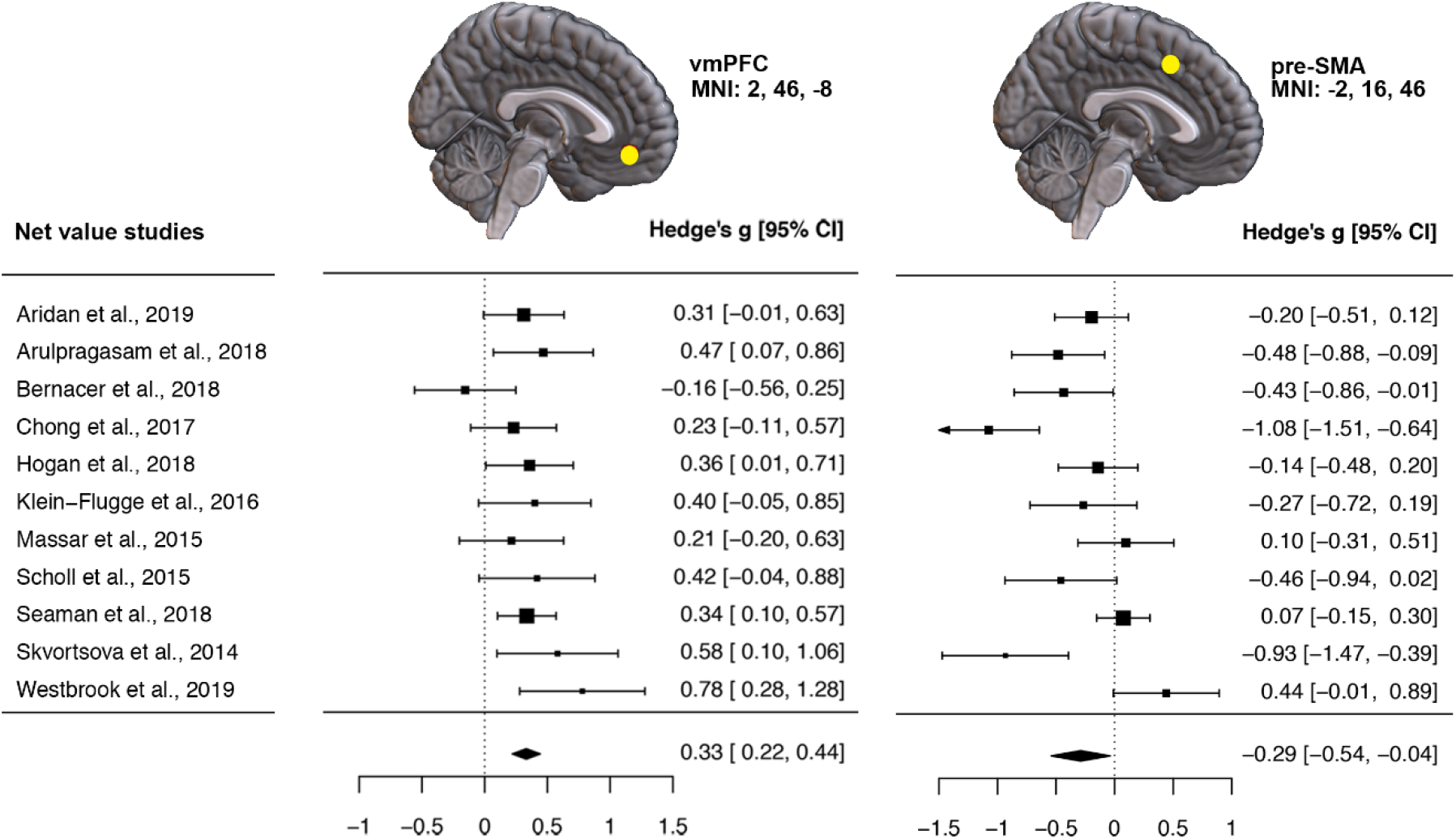
Forest plot illustrating activation related to net value in the vmPFC and pre-SMA ROIs in studies with statistical maps. The vmPFC is activated (Hedge’s g= 0.22, 95% CI [0.22, 0.44]) and the pre-SMA is deactivated (Hedge’s g= −0.28, 95% CI [−0.52, −0.03]) during effort-reward integration.

**Table 2.**
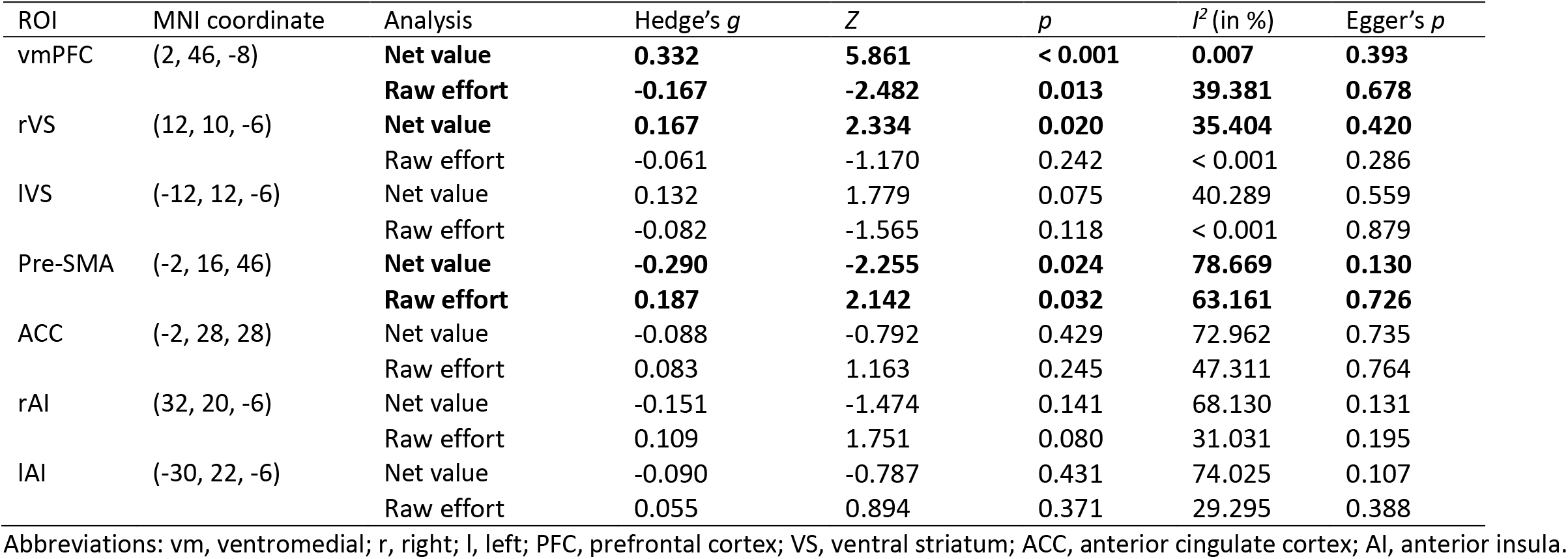
Results of ROI analyses

### 3.2 Whole-Brain Analysis

#### 3.2.1 Prospective Effort

We first examined brain regions that were consistently associated with the valuation of prospective effort demands. As illustrated in Figure 4a, the analysis yielded positive effects clustered in the right pre-SMA and adjacent caudal ACC (see Table 3). At a more lenient, uncorrected *p* < 0.001 threshold, other positive foci were detected in the left SMA, right precuneus, and left middle frontal gyrus, and negative foci were detected in the bilateral vmPFC/OFC and left middle temporal gyrus.

**Figure 4.**
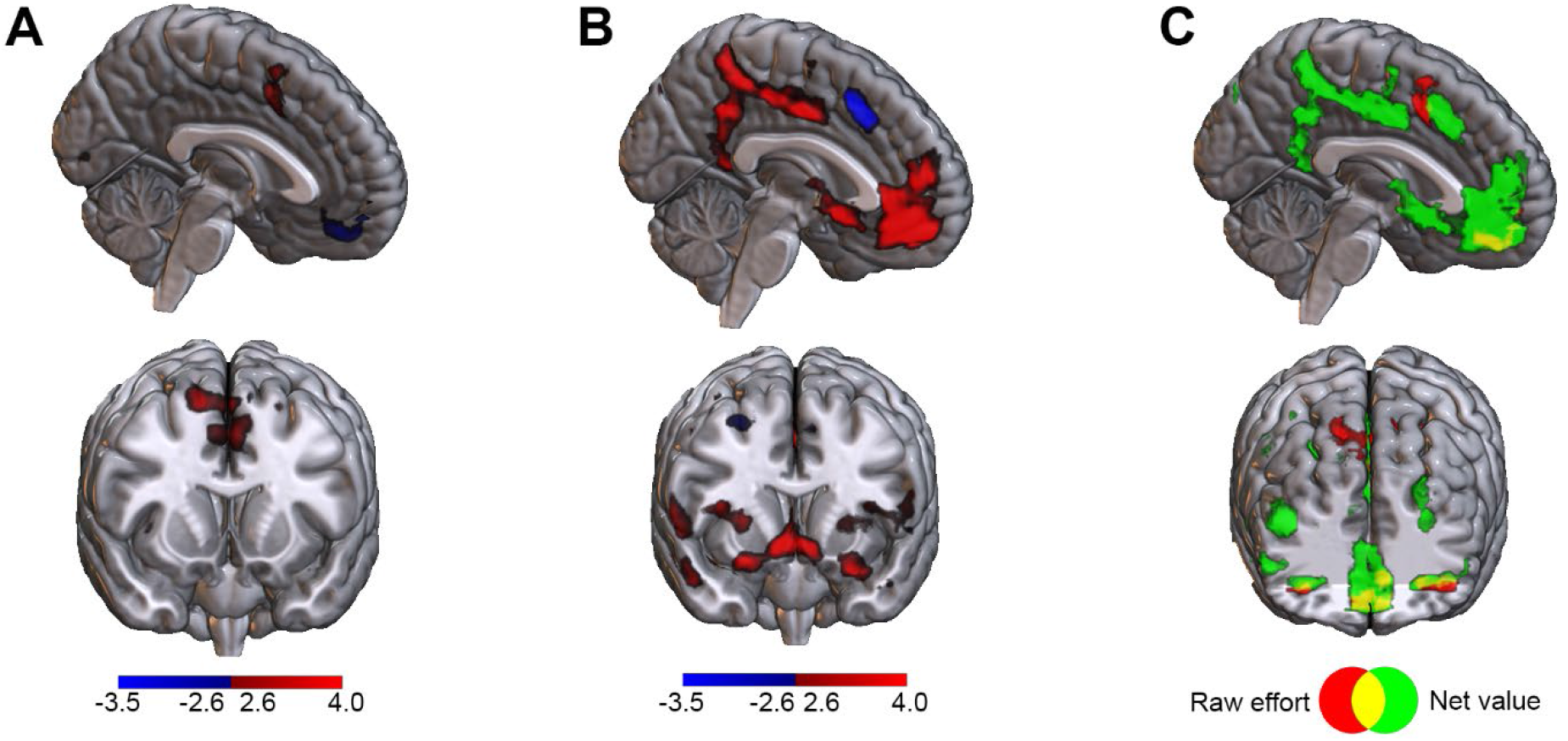
Whole-brain meta-analytic results. A: neural activity related to pure effort cost representation; B: neural activity related to net value; and C: their conjunction based on absolute values. Display threshold: uncorrected *p* < 0.005 at voxel level.

**Table 3.**
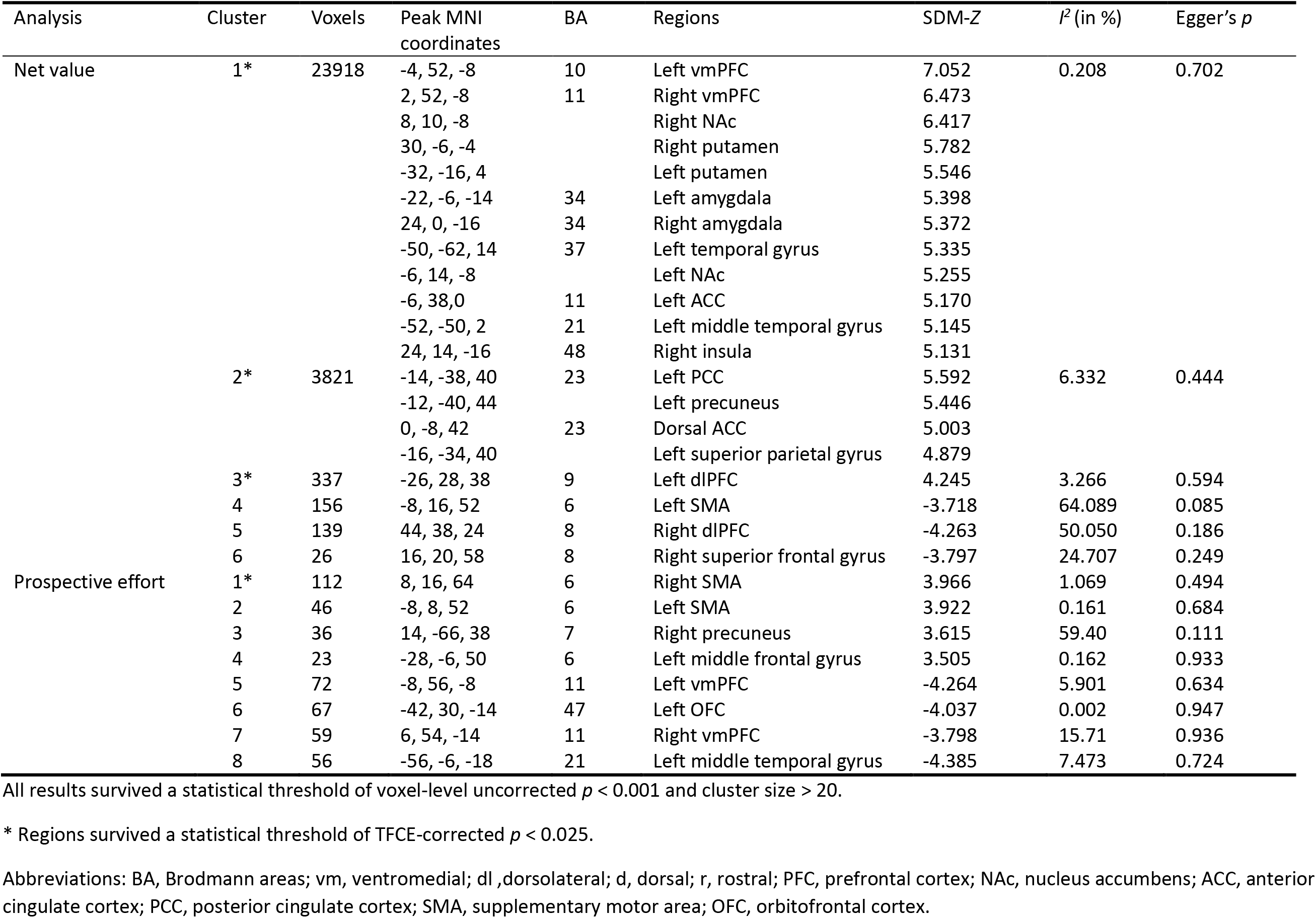
Results of whole-brain analyses

Heterogeneity *I*^*2*^ statistics, funnel plots and Egger regressions did not detect excess heterogeneity or publication bias in any significant clusters in the TFCE-corrected findings. However, in the uncorrected analysis, activation in a cluster in the right precuneus was found to be associated with extreme heterogeneity (*I*^*2*^=59.50%).

#### 3.2.2 Net Value

Next, we examined brain regions that were consistently associated with net value encoding. As illustrated in Figure 4b, the analysis yielded a large cluster connecting cortical and subcortical regions of the medial PFC, VS, dorsal striatum (bilateral putamen and left caudate), and temporal gyrus (see Table 3). Analysis also yielded consistent net value activations in a cluster consisting of the bilateral medial and posterior cingulate cortex and precuneus and a separate cluster in the left middle frontal gyrus. Deactivations were concentrated in small clusters in the left SMA, right dorsolateral PFC (dlPFC), and right superior frontal gyrus, although these deactivations were only detectable at a lenient uncorrected *p* < 0.001 threshold.

In addition, heterogeneity *I*^*2*^ statistics, funnel plots and Egger regressions showed no evidence of excess heterogeneity or publication bias in any of the significant clusters for the main net value or single SV subgroup TFCE-corrected results. No evidence of publication bias was detected in the uncorrected net value analysis, however deactivations in the left SMA and right dlPFC had *I*^*2*^ statistics of 64.09% and 50.05% respectively, suggesting that findings in these two regions were highly heterogenous.

#### 3.2.3 Conjunction Analysis

Finally, we performed a conjunction analysis to identify areas that are sensitive to *both* net value and effort requirements. Due to the exploratory nature of this analysis, we used a lenient threshold of uncorrected p < 0.001 at voxel level and k > 20 at cluster level. Note that we used absolute values in the conjunction analysis because of the clearly dissociable effects found in the main prospective effort and net value meta-analyses. We found that the vmPFC and left lateral orbitofrontal cortex were significantly activated by net value but deactivated by effort requirement. The activation pattern was reversed in the pre-SMA and caudal ACC (Figure 4c). However, all of these findings were not detectable after whole-brain TFCE-correction.

#### 3.2.4 Supplementary analyses

To ensure that the results of the net value meta-analysis were not driven by choice difficulty, we reran our analysis excluding four experiments that used the value of two options as their net value metric (e.g. difference in SV of more vs less effortful option). Importantly, the vmPFC and bilateral VS remained to be the foci with highest effect sizes, and the whole-brain activation pattern was qualitatively similar (see Table S1 and Figure S11), suggesting that our main findings were not influenced by the cognitive demands of comparing two options. Moreover, to ensure that our findings were robust when using a broader definition of net value, we also repeated our analysis including two additional studies that used reward and effort interactions as a measure of net value. Main foci and whole-brain activation patterns remained qualitatively similar to the initial net value meta-analysis (see Table S2 and Figure S12). However, deactivations associated with net value were not detected in these supplementary analyses, suggesting that the deactivations in the SMA detected in the main meta-analysis were not robust.

## 4. Discussion

We conducted a series of combined coordinate- and image-based meta-analyses to examine the neural substrates of effort-based valuation. We first investigated neural activity related to raw effort and net value in seven *a priori* ROIs previously implicated in value-based decision-making. We found these regions could be broadly divided into two groups that exhibited distinct activity pattern during these two processes, with the vmPFC and pre-SMA as the central node of each. Specifically, the vmPFC was consistently activated during net value integration but deactivated for raw effort representation, whereas the pre-SMA displayed the opposite pattern. The exploratory whole-brain and conjunction analyses further corroborate the ROI analyses. These findings provide strong evidence for a dissociable role of the vmPFC and pre-SMA in the valuation of effort costs, and implicate these two regions as core components of a network that drives motivated behavior.

Our findings provide comprehensive evidence that effort-related net value integration is processed in a network centered around the vmPFC and VS. Accumulating evidence implicates the vmPFC as a general hub for value integration, as it has been identified to signal net value of rewards across different cost domains, such as risk and delay (Croxson et al., 2009; Hogan et al., 2019; Kable and Glimcher, 2007; Levy et al., 2010; Peters and Büchel, 2009; Schmidt et al., 2012; Westbrook et al., 2019). Additionally, the network including the vmPFC have been implicated in tracking net values across reward domains (i.e., primary, secondary, and aesthetic rewards), reward processing phases (Bartra et al., 2013; Clithero and Rangel, 2013; Levy and Glimcher, 2012; Sescousse et al., 2013), reward rates, and the value of current and previous offers (Mehta et al., 2019). These findings are therefore consistent with prominent neuroeconomic accounts which propose that the vmPFC represent the net value of an option in a ‘common currency’, in order to facilitate value comparison during decision-making (Padoa-Schioppa, 2011; Rangel et al., 2008; Westbrook and Braver, 2015).

One would hypothesize that a region involved in representing net value would also scale with effort demands. Except for the vmPFC, our study did not find that other net-value-related regions, such as the VS, meet this requirement. These findings are at odds with previous reports that the VS signals prospective effort costs in humans, both in the presence (Westbrook et al., 2019) and absence (Suzuki et al., 2020) of reward information. Moreover, dorsal parts of the striatum have also been found to track both effort costs (Burke et al., 2013; Guitart-Masip et al., 2012; Klein-Flugge et al., 2016; Kurniawan et al., 2010, 2013; Yang et al., 2016) and net value of prospective effortful rewards (Klein-Flugge et al., 2016; Seaman et al., 2018). However, our results implicate motor-related regions of the striatum, particularly the putamen, as signaling net value alone. One plausible explanation is that the striatum signals both net value and prospective effort during this time window, but that the simultaneous nature of these signals inhibits detection. Studies that have experimentally isolated prospective effort demands from net value, however, did not find that the striatum was activated by effort alone (Arulpragasam et al., 2018), leaving role of the striatum in effort anticipation as a salient question for future investigation.

Finally, both main and supplementary analyses consistently identified a variety of parietotemporal regions as scaling positively and uniquely with net value representations. While these regions (i.e. intraparietal lobule, intraparietal sulcus, temporal pole, etc.) have been previously implicated in SV encoding of effortful rewards (Chong et al., 2017; Massar et al., 2015), they also play a critical role in perceptual decision-making (Keuken et al., 2014), attention (Husain, 2019), risk weighting (Mohr et al., 2010), and decision difficulty (Westbrook et al., 2019). Their notable absence in reward processing (Keuken et al., 2014; Sescousse et al., 2013) may thus suggest that these parietotemporal regions are involved in high-level perceptual and cognitive functions associated with task demands as opposed to net value computation.

Previous studies have identified effort-related net value signals in other frontal regions, such as the pre-SMA and ACC, which suggests that these regions may be specifically relevant for effort-reward integration. In the current meta-analysis, however, we found that these regions – in particular, the pre-SMA and adjacent caudal ACC – all scaled positively with raw effort costs and, albeit less robustly, scaled negatively with net value. Such a pattern suggests that these regions are more likely to be involved in the processing of effort-related costs, rather than value integration per se. These findings align closely with a previous transcranial magnetic stimulation study, in which disruption of the SMA led to decreased effort perception (Zénon et al., 2015). The pre-SMA and dorsal ACC are also recruited to process other types of costs, such as risk (Mohr et al., 2010) and delay (Schüller et al., 2019). A plausible mechanism, therefore, is that these regions serve as a domain-general hub for cost encoding and transfer the cost information to the vmPFC for calculation of net value. Alternatively, neuroeconomic models of effort-based decision-making have posited that the ACC, in particular, is involved in good-to-action transformation (Padoa-Schioppa, 2011). Thus, another plausible mechanism is that the vmPFC computes and compares the net value of separate options and passes choice preference to action selection regions, such as the pre-SMA and ACC, for conversion to motor output.

Despite strong evidence about the involvement of the caudal ACC, which is close to the pre-SMA, in effort costs processing, it should be noted that the ACC, as a whole, is highly heterogeneous (Neubert et al., 2015; Yu et al., 2011). Indeed, the whole-brain results showed distinct activation patterns across the ACC, in which net-value-related activation was mainly observed in the ventral ACC, whereas cost-related activation in the dorsal ACC. These findings suggest that subregions of the ACC could be linked to different aspects of the effort-related valuation, which may also partly explain the fact that some studies identified net value signals in the ACC (Klein-Flugge et al., 2016; Massar et al., 2015). Moreover, net-value-related activation may emerge in the dorsal ACC if it is highly correlated with other confounding variables, such as decision difficulty (Shenhav et al., 2013). It is particularly plausible for studies that have used the SV difference between two options as the net value parameter, as it often approximates decision difficulty (Chong et al., 2017; Klein-Flugge et al., 2016). Notably, studies that have experimentally isolated net value and decision difficulty showed that the cognitive control network, including the dorsal ACC and other frontoparietal regions, tracked the latter but not the former (Hogan et al., 2019; Westbrook et al., 2019).

The current study has some limitations. First, the sample size of the net value analysis is relatively small. Although the inclusion of statistical images partly offsets this issue, the number of included studies still limited our ability to further explore the effects of potential moderators, such as effort type (i.e., physical vs. cognitive), parameter type (i.e., difference in SV vs. SV of one option), effort execution requirement (i.e., real vs. hypothetical), and reward probability (i.e., cumulative vs. random payout). Because effort-based decision-making is sensitive to reward probability (Barch et al., 2014; Soder et al., 2020; Treadway et al., 2012) and opportunity costs (Otto and Daw, 2019), future research should directly explore the interaction between effort demand and other cost domains and/or task features. Second, the majority of the included studies focused on physical effort measured by handgrip devices. These findings should be treated cautiously when generalizing to other formats of effort. Finally, the meta-analytic results reflected consistent regional activations across studies. Although our study identified critical brain regions related to effort-related value integration or cost encoding, how these regions interact with each other to achieve the dynamic valuation process remains to be elucidated by studies using task-based connectivity technique (Hauser et al., 2017) or imaging methods with higher temporal resolution (e.g., magnetoencephalography).

In conclusion, this study is the first to use combined image- and coordinate-based meta-analyses to examine neural activity related to effort-related costs and net value. The results showed the pre-SMA is involved in cost representation of prospective effort independent of rewards. In contrast, the vmPFC and VS, which have been implicated in value integration in other cost domains, are also involved in effort-reward integration. These findings further clarify the neural mechanisms underlying effort-related valuation and may provide candidate intervention targets for patients with decreased motivation to exert effort to obtain rewards.

## Acknowledgement

PL-G was supported by a fellowship from “la Caixa” Foundation (LCF/BQ/DI19/11730047). Y-WY was supported by the PhD fellowship of the Einstein Center for Neurosciences Berlin. TT-JC was supported by the Australian Research Council (DP 180102383 and DE 180100389). The authors would like to thank Nadav Aridan, Amanda Arulpragasam, Javier Bernacer, Michael Chee, Vikram Chib, Claudie Gaillard, Erica Grodin, Tobias Hauser, Masud Husain, Miriam Klein-Flügge, Irma Kurniawan, Stijn Massar, Il Ho Park, Mathias Pessiglione, Ceyda Sayalı, Jacqueline Scholl, Kendra Seaman, Vasilisa Skvortsova, Michael Treadway, and Andrew Westbrook for sharing whole-brain statistical maps or peak coordinates.

