## Supplementary Materials for "The neural basis of effort valuation: A meta-analysis of functional magnetic resonance imaging studies"

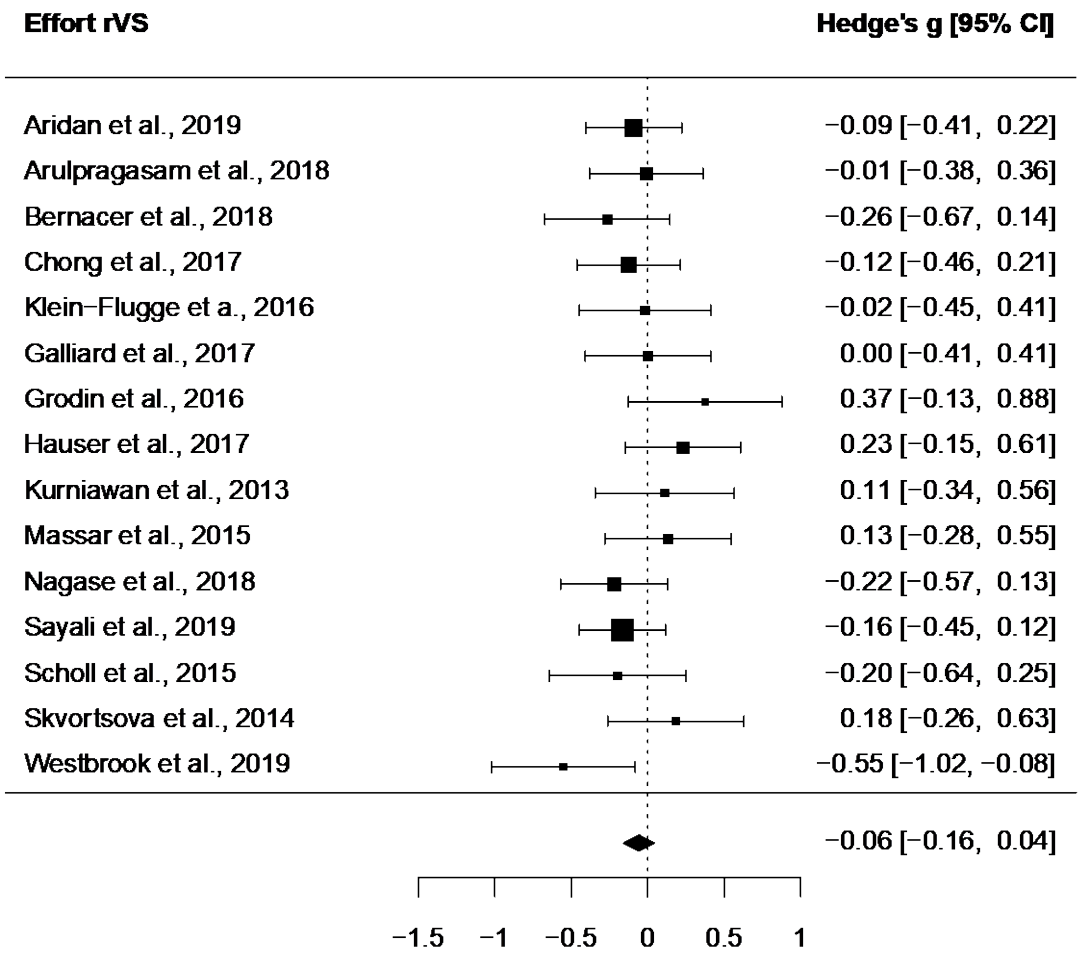


Figure S1. Forest plot illustrating activation related to raw effort in the right VS ROI in studies with statistical maps.

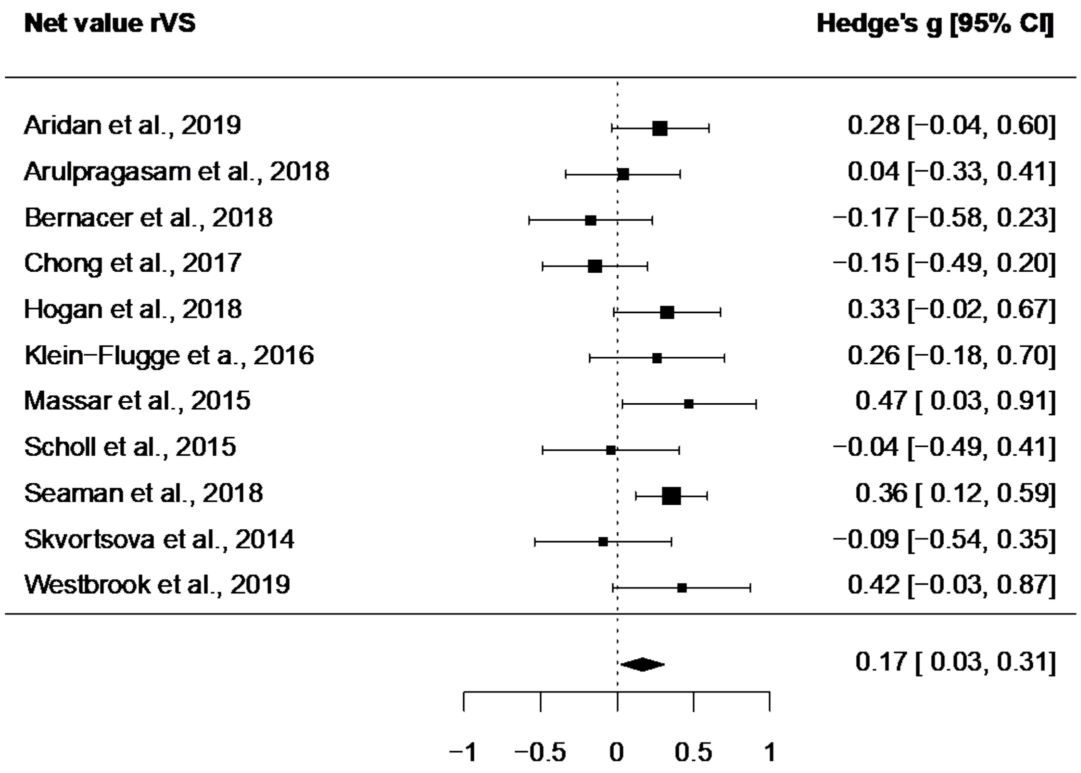


Figure S2. Forest plot illustrating activation related to net value in the right VS ROI in studies with statistical maps.

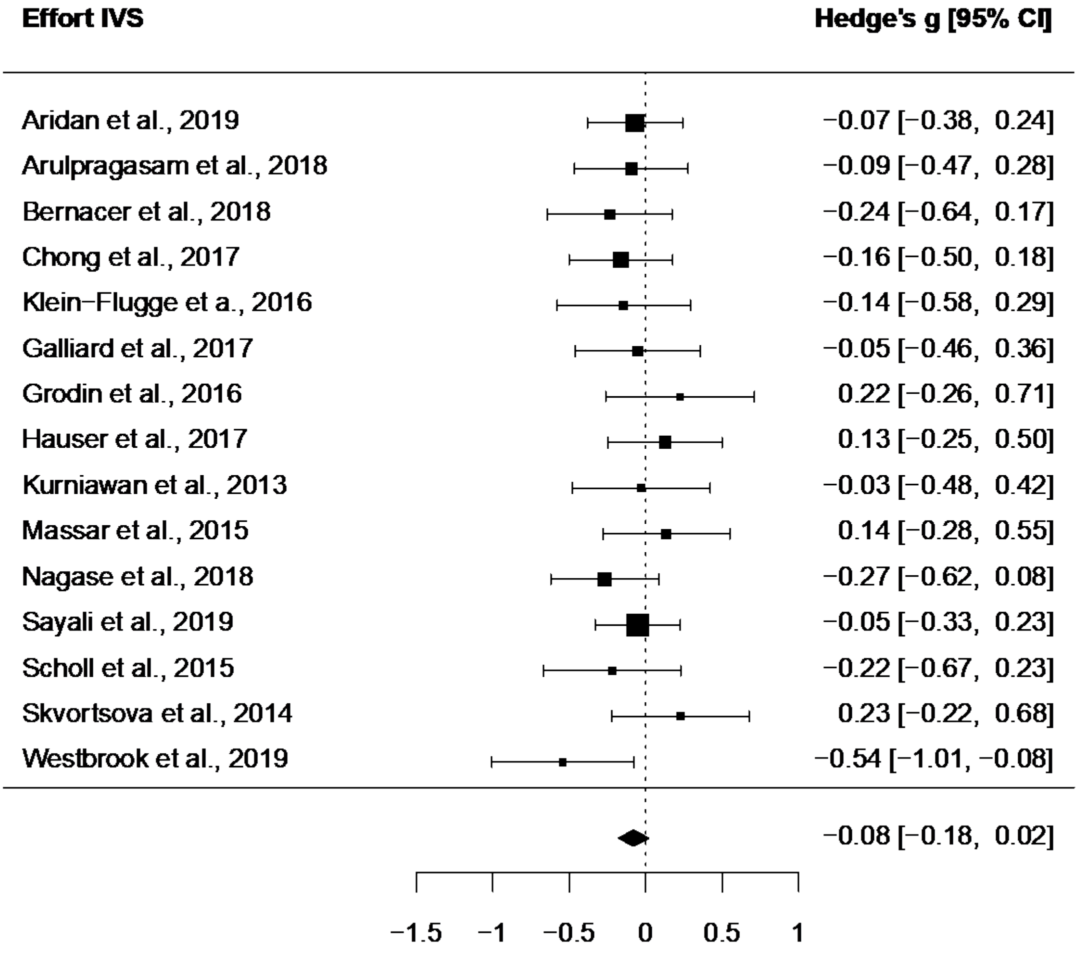


Figure S3. Forest plot illustrating activation related to raw effort in the left VS ROI in studies with statistical maps.

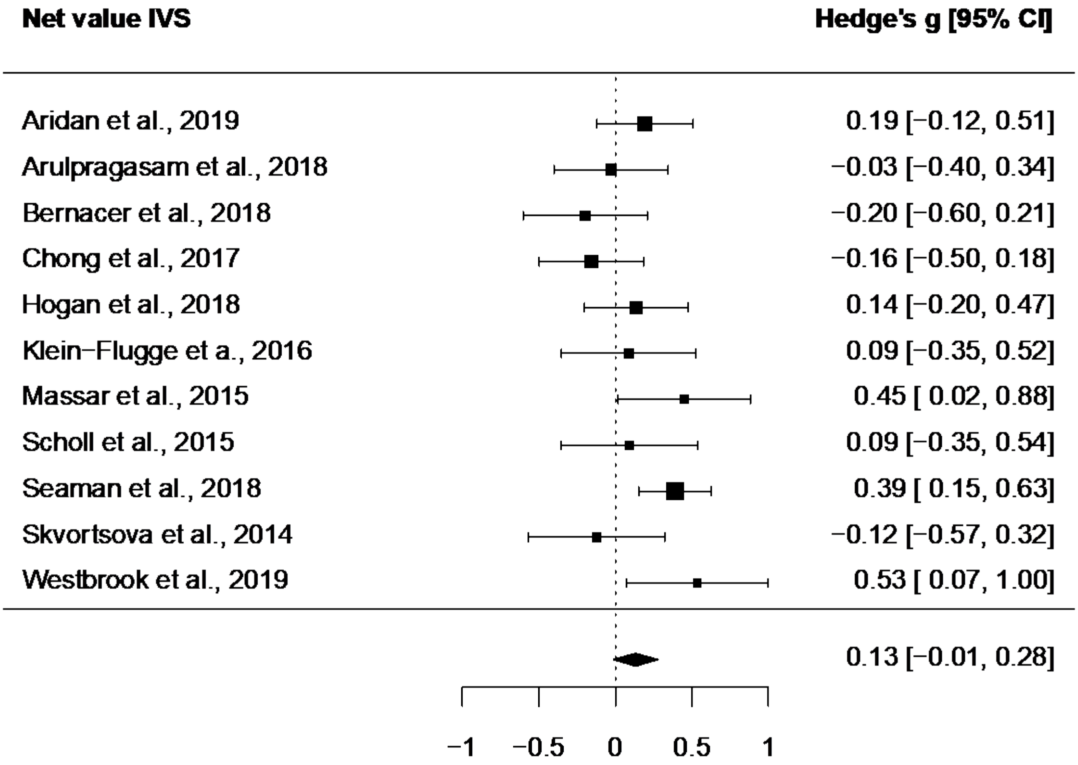


Figure S4. Forest plot illustrating activation related to net value in the left VS ROI in studies with statistical maps.

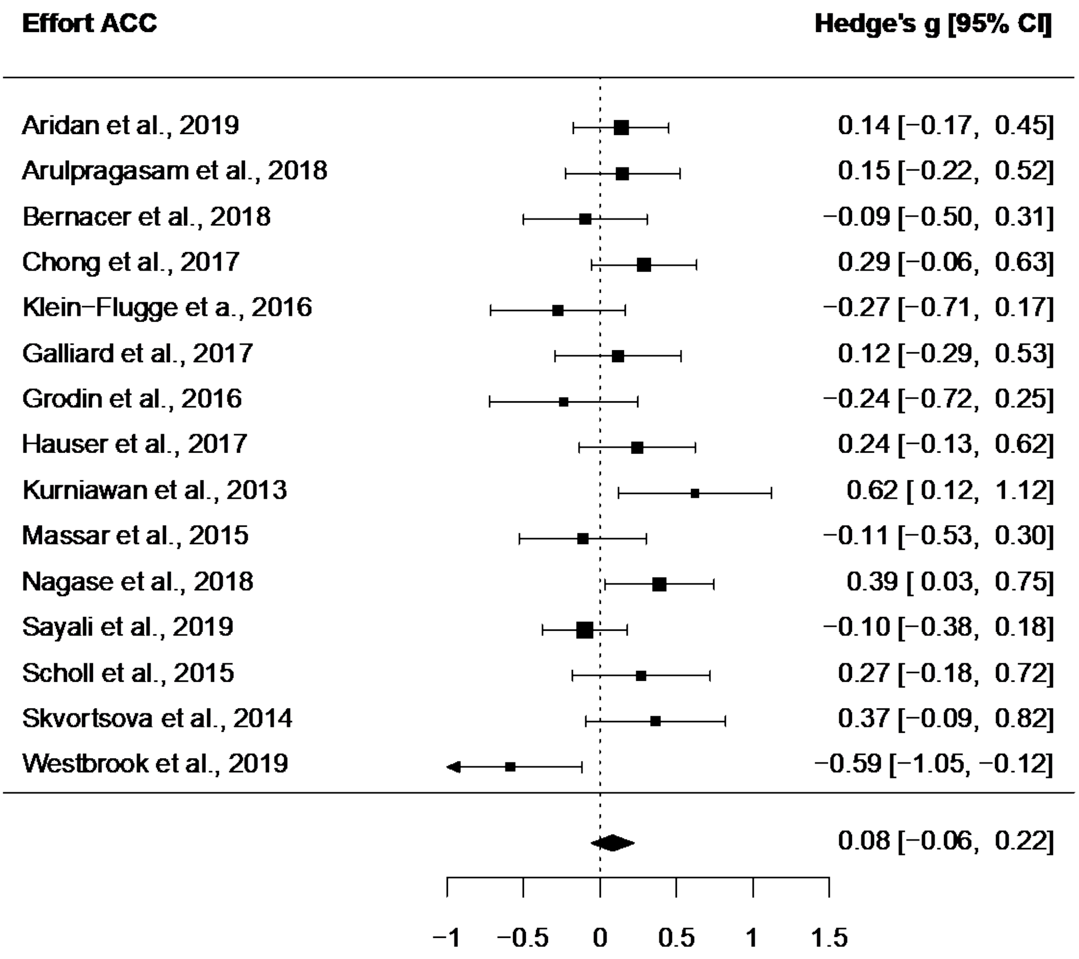


Figure S5. Forest plot illustrating activation related to raw effort in the ACC ROI in studies with statistical maps.

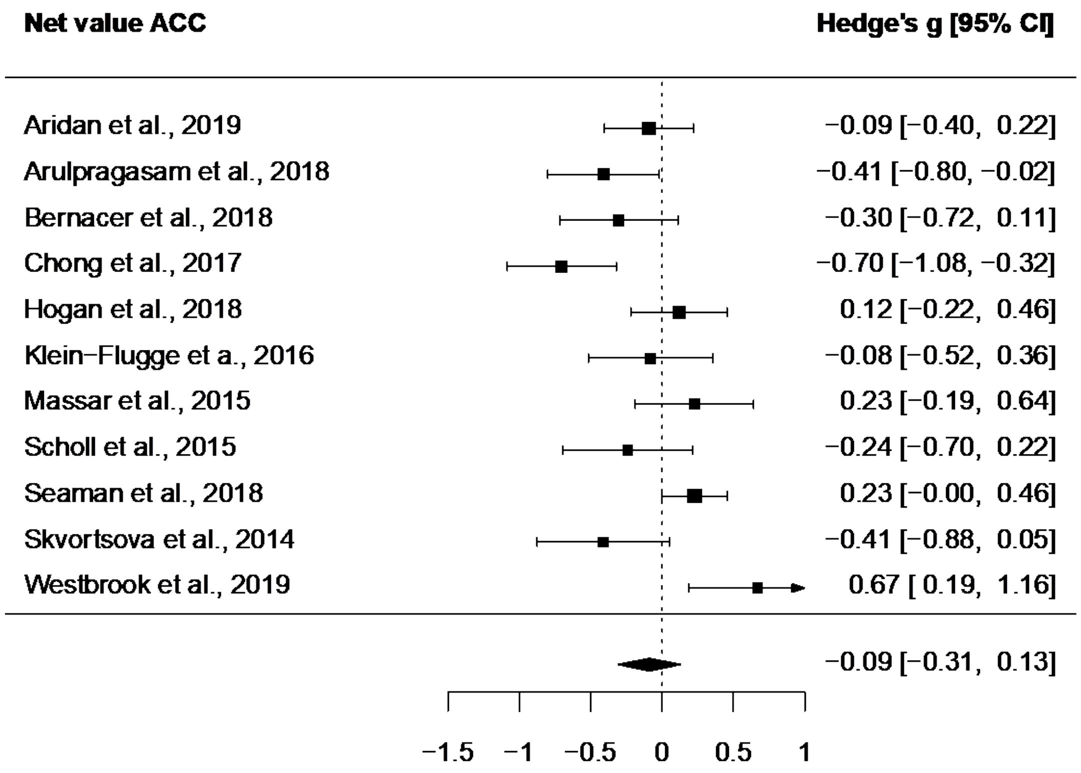


Figure S6. Forest plot illustrating activation related to net value in the ACC ROI in studies with statistical maps.

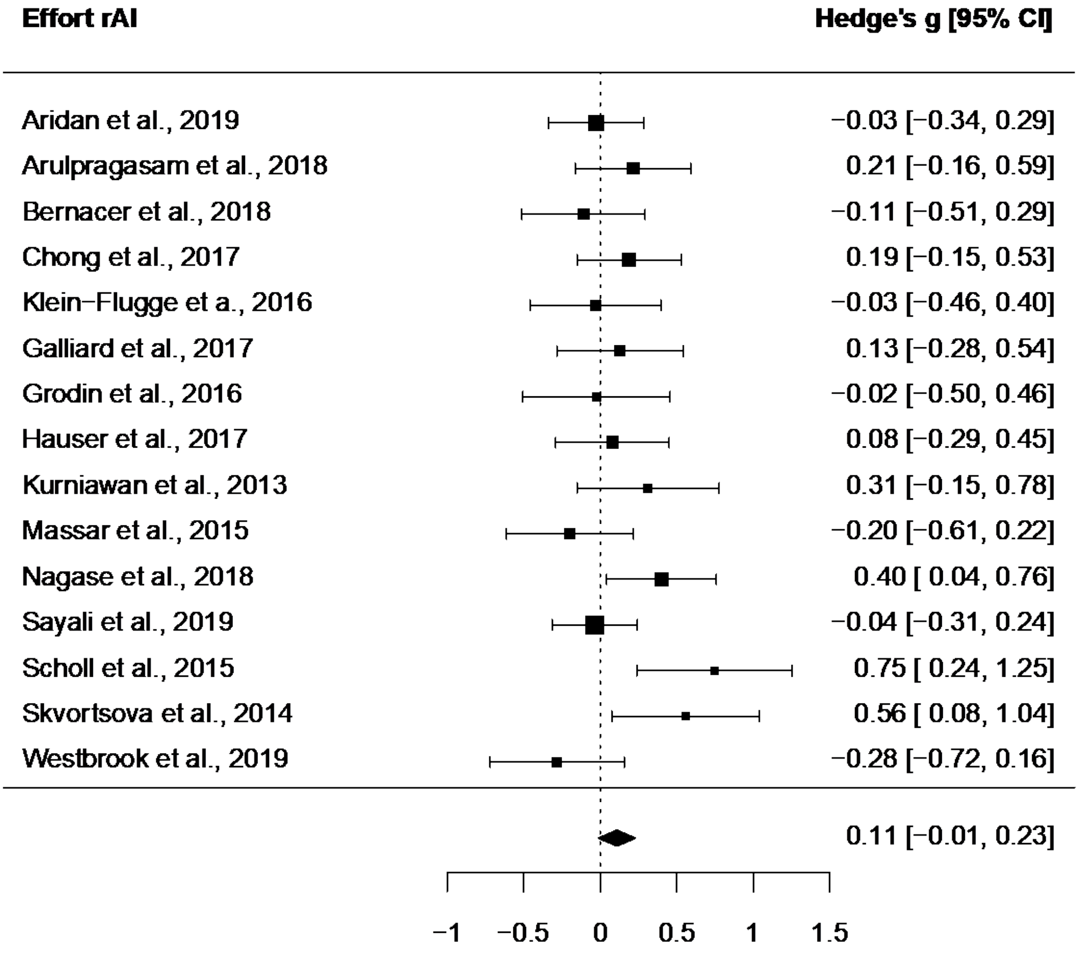


Figure S7. Forest plot illustrating activation related to raw effort in the right AI ROI in studies with statistical maps.

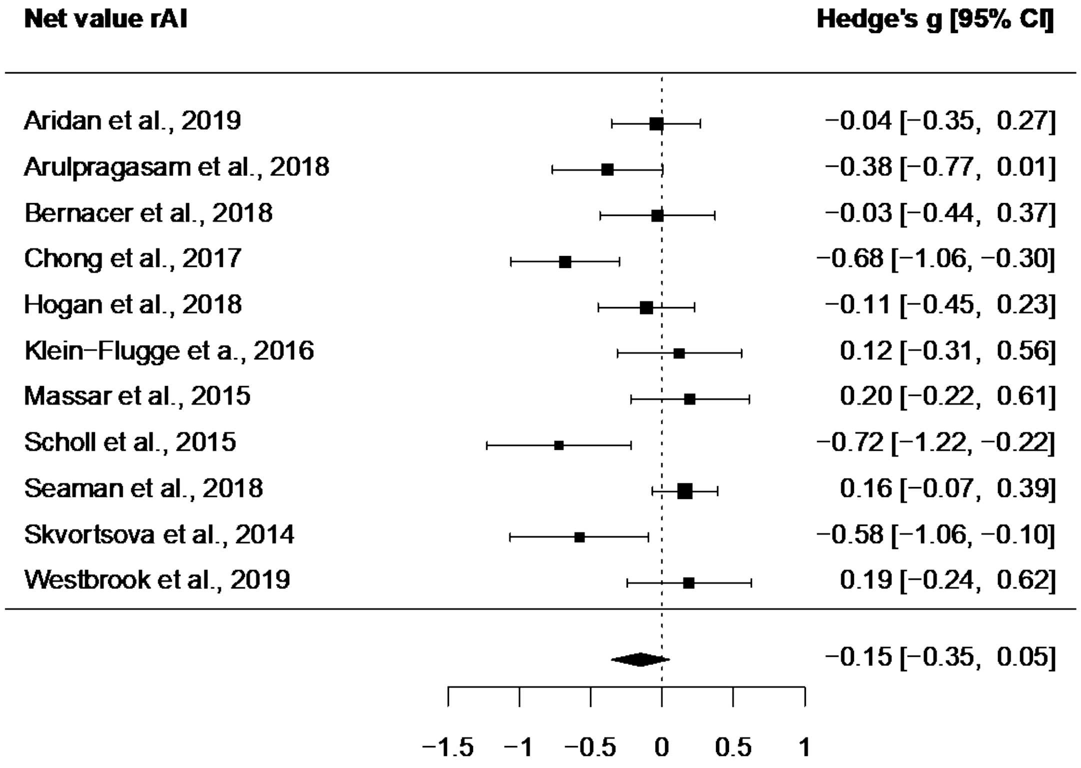


Figure S8. Forest plot illustrating activation related to net value in the right AI ROI in studies with statistical maps.

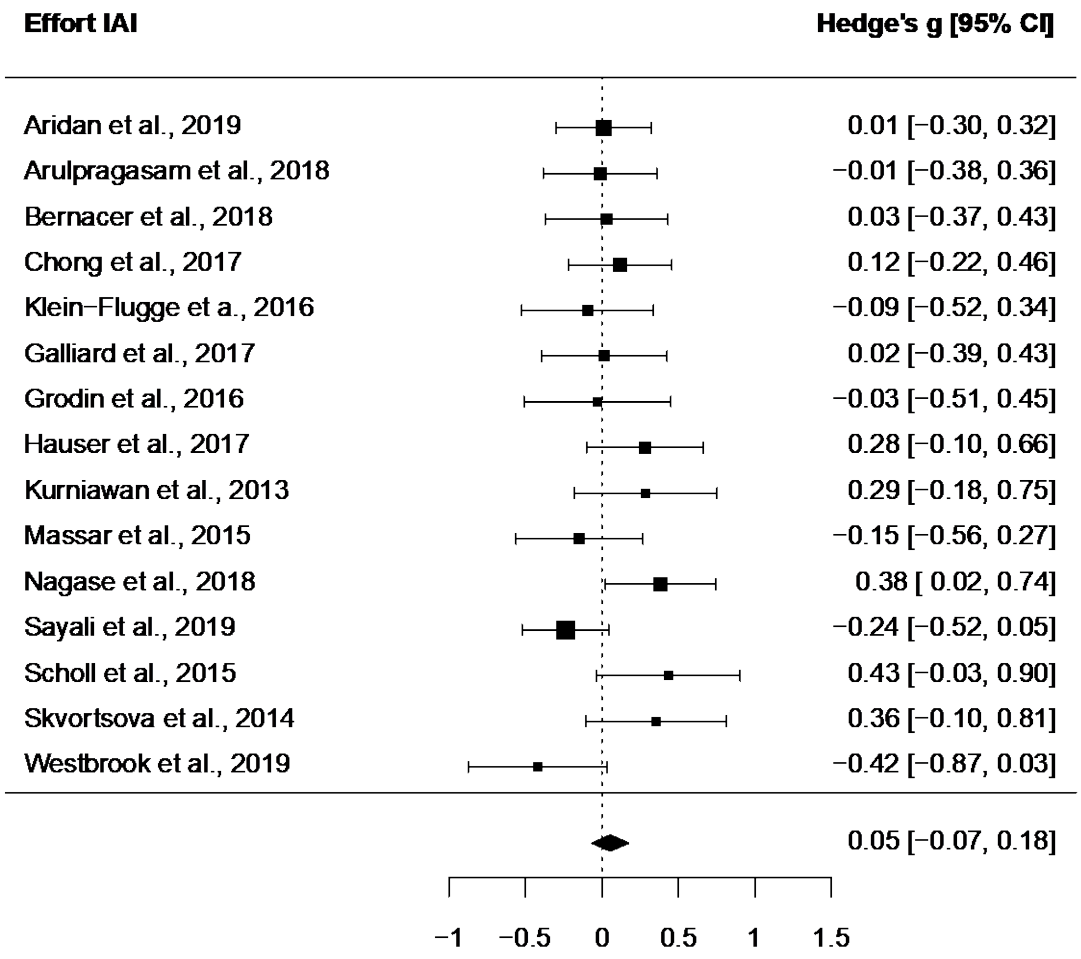


Figure S9. Forest plot illustrating activation related to raw effort in the left AI ROI in studies with statistical maps.

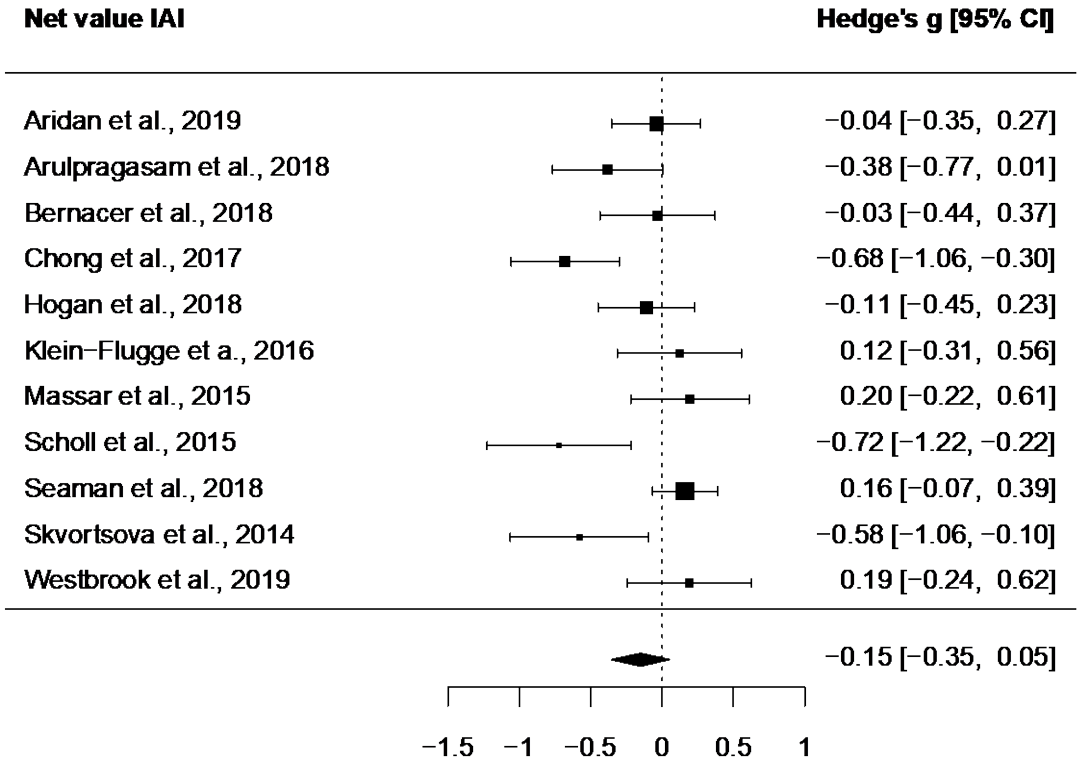


Figure S10. Forest plot illustrating activation related to net value in the left AI ROI in studies with statistical maps.

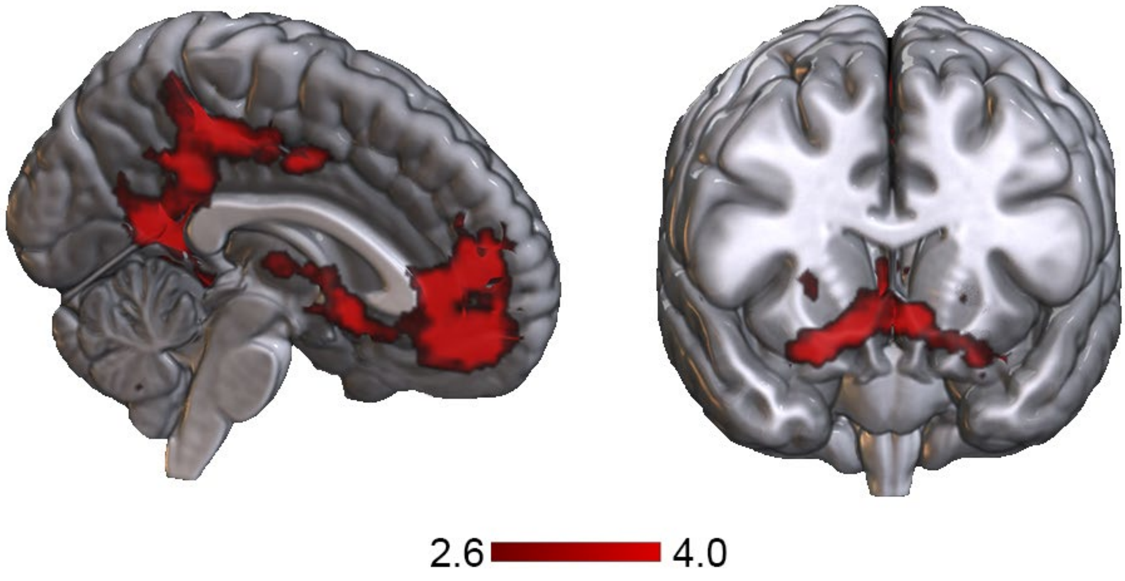


Figure S11. Whole-brain meta-analytic results of single option net value subgroup analysis. Results use an uncorrected p < 0.001 threshold and represent neural activity consistently related to effort-reward integration in studies using parameters that only include the net value of one choice option (N=11). Findings generally replicate activation activity in the main net value meta-analysis but did not detect any consistent deactivations.

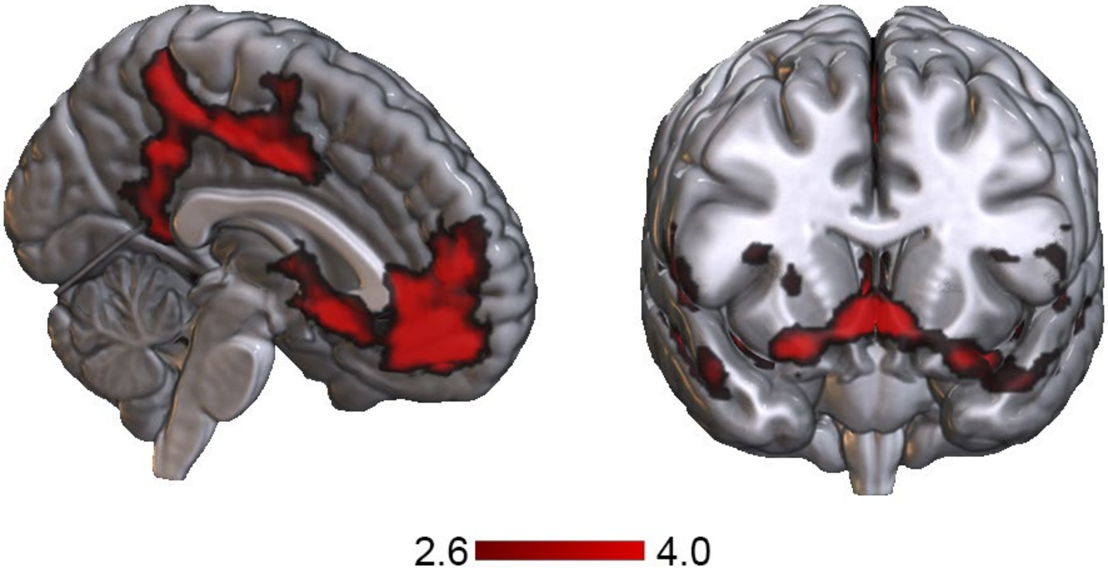


Figure S12. Whole-brain meta-analytic results of net value supplementary analysis including studies with EffortXReward interaction parameters (N=17). Results use an uncorrected $p < 0.001$ threshold. Findings generally replicate activation activity in the main net value meta-analysis but did not detect any consistent deactivations.

Table S1. Effect of 1 Parameter Net Value on BOLD

| Cluster | Voxels | Peak MNI coordinates | BA | Regions | SDM-Z | $I^2$ | Egger's *p* |
|---|---|---|---|---|---|---|---|
| 1* | 29860 | -6, 52, -6 | 10 | Left vmPFC | 7.152 | 0.660 | 0.793 |
| | | 4, 48, -8 | 10 | Right vmPFC | 6.638 | | |
| | | 8, 10, -8 | | Right NAc | 6.178 | | |
| | | -8, 12, -8 | | Left NAc | 5.575 | | |
| | | -24, -2, -16 | 34 | Left amygdala | 5.365 | | |
| | | 24, 0, -16 | 34 | Right amygdala | 5.477 | | |
| | | -4, 12, -6 | 25 | Left NAc | 5.301 | | |
| | | 8, 52, 12 | 10 | dmPFC | 5.212 | | |
| | | -6, 38, 0 | 11 | Left rACC | 5.172 | | |
| | | -26, 36, -10 | 11 | Left lateral OFC | 5.093 | | |
| | | -10, -48, 8 | | Left PCC | 4.951 | | |
| | | 32, -12, 2 | 48 | Left putamen | 4.835 | | |
| 2* | 344 | -24, 26, 40 | 9 | dlPFC | 5.116 | 0.148 | 0.755 |
| 3* | 76 | 44, 40, 2 | 45 | Right inferior frontal gyrus | 3.446 | 0.107 | 0.847 |
| 4* | 50 | -24, -38, 60 | 3 | Left postcentral gyrus | 3.263 | 1.336 | 0.402 |
| 5 | 55 | 44, -54, 14 | 21 | Right posterior temporal gyrus | 3.844 | 0.427 | 0.544 |

Note: Data from 11 studies were included in this analysis. All results survived a statistical threshold of voxel-level uncorrected $p < 0.001$ and cluster size > 20.

* Regions survived FWER-correction with a TFCE threshold of p < 0.025.

Abbreviations: BA, Brodmann areas; vm, ventromedial; dm, dorsomedial; dl, dorsolateral; r, rostral; PFC, prefrontal cortex; NAc, neural accumbens; ACC, anterior cingulate cortex; PCC, posterior cingulate cortex; OFC, orbitofrontal cortex.

Table S2. Effect of Net Value (with Effort X Reward interaction) on BOLD

| Cluster | Voxels | Peak MNI coordinates | BA | Regions | SDM-Z | $I^2$ | Egger's *p* |
|---|---|---|---|---|---|---|---|
| 1* | 24611 | -4, 52, -8 | 10 | Left vmPFC | 6.970 | 0.312 | 0.792 |
| | | 2, 50, -8 | 10 | Right vmPFC | 6.625 | | |
| | | 8, 10, -8 | | Right NAc | 6.482 | | |
| | | 32, -12, 4 | 48 | Right putamen | 5.814 | | |
| | | -2, 40, -10 | 11 | Left rACC | 5.805 | | |
| | | 10, 50, 12 | 32 | Right dmPFC | 5.787 | | |
| | | -6, 14, -8 | 25 | Left caudate | 5.172 | | |
| | | 24, 0, -16 | 34 | Right amygdala | 5.543 | | |
| | | -22, -6, -14 | 34 | Left amygdala | 5.528 | | |
| | | -32, -16, 4 | 48 | Left insula | 5.493 | | |
| | | 34, 34, 12 | 47 | Left lateral OFC | 4.932 | | |
| 2* | 4473 | -14, -38, 40 | | Left PCC | 5.547 | 5.050 | 0.513 |
| | | -12, -40, 44 | | Left precuneus | 5.411 | | |
| | | 0, -8, 42 | 23 | ACC | 5.065 | | |
| | | 8, -34, 52 | | Right PCC | 4.952 | | |
| 3* | 520 | -26, 28, 38 | 9 | Left dlPFC | 4.200 | 2.892 | 0.622 |
| 4 | 130 | 44, 38, 24 | 45 | Right inferior frontal gyrus | -4.173 | 48.183 | 0.255 |
| 5 | 126 | -8, 16, 52 | 6 | Left SMA | -3.612 | 62.458 | 0.130 |
| 6 | 26 | 16, 20, 58 | 8 | Right dlPFC | -3.635 | 35.232 | 0.403 |

Note: Data from 17 studies were included in this analysis. All results survived a statistical threshold of voxel-level uncorrected $p < 0.001$ and cluster size > 20.

* Regions survived FWER-correction with a TFCE threshold of $p < 0.025$.

Abbreviations: BA, Brodmann areas; vm, ventromedial; dm, dorsomedial; dl, dorsolateral; r, rostral; PFC, prefrontal cortex; NAc, neural accumbens; ACC, anterior cingulate cortex; PCC, posterior cingulate cortex; SMA, sensory motor area.
